# Mechanism of genome interrogation: How CRISPR RNA-guided Cas9 proteins locate specific targets on DNA

**DOI:** 10.1101/163501

**Authors:** A. A. Shvets, A. B. Kolomeisky

## Abstract

The ability to precisely edit and modify genome opens endless opportunities to investigate fundamental properties of living systems as well as to advance various medical techniques and bioengineering applications. This possibility is now close to reality due to a recent discovery of the adaptive bacterial immune system, which is based on clustered regularly interspaced short palindromic repeats (CRISPR)-associated proteins (Cas) that utilize RNA to find and cut the double-stranded DNA molecules at specific locations. Here we develop a quantitative theoretical approach to analyze the mechanism of target search on DNA by CRISPR RNA-guided Cas9 proteins, which is followed by a selective cleavage of nucleic acids. It is based on a discrete-state stochastic model that takes into account the most relevant physical-chemical processes in the system. Using a method of first-passage processes, a full dynamic description of the target search is presented. It is found that the location of specific sites on DNA by CRISPR Cas9 proteins is governed by binding first to protospacer adjacent motif (PAM) sequences on DNA, which is followed by reversible transitions into DNA interrogation states. In addition, the search dynamics is strongly influenced by the off-target cutting. Our theoretical calculations allows us to explain the experimental observations and to give experimentally testable predictions. Thus, the presented theoretical model clarifies some molecular aspects of the genome interrogation by CRISPR RNA-guided Cas9 proteins.

Insert Received for publication Date and in final form Date.

## 1 INTRODUCTION

One of the most surprising recent discoveries in biology is the finding that many bacteria have an RNA-supported adaptive immune system, which very efficiently targets and eliminates any outside genetic material (1–4). Responding to the invasion of viruses and plasmids, bacteria integrates short fragments of foreign nucleic acids into its own genome in a special region with a repetitive pattern, which is known as a CRISPR (clustered regularly interspaced short palindromic repeat) (2, 4). The CRISPR region consists of two elements: repeat segments (approximately 20-50 base pairs) that have the same composition in the given bacteria, and the spacers of similar length that are unique and represent the segments of foreign nucleic acids. Transcription of the CRISPR region produces short RNA molecules, which are called CRISPR-derived RNA (crRNA), that contain sequences complimentary to previously encountered foreign nucleic acids. These crRNA then direct CRISPR-associated (Cas) proteins to find and destroy the complimentary target sequences on invading viral or plasmid DNA molecules by cutting them in pieces (2, 3). Experiments show that CRISPR-Cas system is a surprisingly simple, powerful and versatile tool for genetic alterations and modifications in various cell types and organisms (5–16). Essentially, CRISPR-Cas systems led to a revolutionary new approach of genome editing with multiple medical and bioengineering applications. But despite these huge technological advances, the fundamental mechanisms of how the CRISPR RNA-guided Cas proteins locate specific targets remains a mystery (5, 6, 12).

There are three major CRISPR-Cas systems in bacteria and archaea that employ slightly different molecular mechanisms to locate and remove foreign nucleic acids material (2, 12). But the simplest of them utilizes only a single protein, CRISPRCas9, for both RNA-guided DNA recognition and for the cleavage. For this reason, this system became very popular for investigations on the mechanisms of CRISPR-associated phenomena and for various genetic manipulations (5, 6, 12, 15, 16). Extensive single-molecule and bulk biochemical measurements (both *in vitro* and *in vivo*) determined that the process of finding the specific sequences on DNA by RNA-guided Cas9 proteins did not involve sliding along the DNA chain but rather multiple short-time collisions (5, 6). At the same time, it was found that both binding and cleavage of DNA by Cas9-RNA complexes require first a recognition of a very short sequence (three nucleotides) known as a protospacer adjacent motif (PAM) (5, 6, 12). These are the sequences that surround the nucleic acid segments from the foreign organisms that are incorporated into the genome of the bacteria in the CRISPR region. This event serves as a signal that the correct sequence might be found next to the PAM segment. Experiments also show that the sequences fully complementary to the guide RNA but without neighboring PAM sequences are ignored by Cas9-RNA complexes (5). In addition, it was also found that the interactions with PAM trigger the catalytic nuclease activity of Cas9 proteins (5, 6, 12). After the PAM recognition, the Cas9-RNA molecule must destabilize the adjacent DNA duplex and initiate the strand separation. It is followed then by the pairing between the target DNA sequence and the crRNA segment, after which the cleavage process starts. Experiments indicate that the formation of RNA-DNA heteroduplex is initiated at the PAM and proceeds in a sequential manner along the target sequence (5, 6).

However, several surprising observations, which could not be easily explained, have been reported in experiments on the CRISPR-Cas9 system (5, 6). It was found that in *in vitro* conditions Cas9-RNA complex does not follow the expected Michaelis-Menten kinetics, and it can be viewed essentially as a single-turnover enzyme (5). The Cas9-RNA complex was also able to locate the correct sequence on DNA quite fast (in less than few minutes) despite the fact that it did not utilize a facilitated diffusion, i.e., a combination of three-dimensional and one-dimensional diffusion motions, to accelerate the search as done by many other proteins, e.g., transcription factors (18–24). Furthermore, there is a non-negligible fraction of events when the Cas9-RNA complex cuts the DNA chain at the wrong sequence (5, 6, 12). This so-called off-target cutting presents a serious challenge for the practical use of CRISPR-associated methods while the mechanism of this phenomenon remains not fully understood (11, 15, 17).

Despite the tremendous importance of CRISPR-associated processes, there is a surprisingly small number of theoretical investigations concerned with molecular mechanisms and dynamics of underlying processes in these systems (13, 27, 28). A recent computational study probed structural dynamics of the CRISPR-Cas9 RNA guided DNA cleavage by utilizing various high-resolution structures of Cas9 proteins in different states (13). It clarified many structural aspects of the process, including the recognition of the PAMs, and the formation of heteroduplex between DNA and RNA segments. However, the computational approach used a very simplified model of the protein (elastic network model) with a limited normal mode analysis. A different approach has been used to develop a systems biology model of CRISPR-Cas9 activities (27). Utilizing methods of statistical thermodynamics and chemical kinetics, a comprehensive model, which takes into account most biochemical and biophysical processes in the system was built. Several important insights on the mechanisms of CRISPR-Cas9 system, including the suggestion that DNA supercoiling controls Cas9 binding and the quantification of the off-target binding frequencies, have been presented. However, there are several problems with this approach. A chemical equilibrium for binding of Cas9-RNA complexes to DNA molecules has been assumed for the process that seems to be very far from equilibrium. In addition, unrealistically large values of the diffusion rates for Cas9 proteins were utilized in calculations. Furthermore, a large number of kinetic parameters were fitted with a limited amount of quantitative data, raising questions on the robustness of the analysis.

In this article, we develop a minimalist theoretical model to describe the target search dynamics of RNA-guided CRISPRCas9 proteins. It is stimulated by a discrete-state stochastic framework that uses a method of first-passage time probabilities for calculating explicitly dynamic properties and takes into account the most relevant biochemical and biophysical processes in the system (24, 29, 30, 32). This framework has been successfully applied before to analyze complex processes in a variety of systems, including protein search on the heterogeneous DNA (29), the effect of DNA looping in the target search of multi-site proteins (30), investigation of the role of conformational transitions in the protein search (31), the mechanism of homology search by RecA protein filaments (32), and the contribution of the intersegment transfer in the protein target search of zincfinger proteins (23). The advantage of this approach is that all results can be obtained analytically, yielding a full dynamic description of the target search process, so that the molecular mechanisms of underlying phenomena can be analyzed. Our calculations predict that the search dynamics by CRISPR-Cas9 proteins is fully controlled by associating/dissociating from the special PAM sites, reversible transitioning into the DNA interrogation sites and by off-target cutting. Furthermore, we explain the single-turnover enzyme observations for Cas9-RNA complex in *in vitro* studies as an effective chemical equilibrium due to very slow rate of dissociation of the nuclease complex after DNA cutting, which quantitatively agrees with experiments.

## Materials and Methods

### Theoretical Model

To describe the genomic interrogation by CRISPR-Cas9 proteins, we propose a discrete-state stochastic model as presented in Fig. 1. Based on experimental observations (5, 6, 12), it is suggested that the following sequence of events is taking place in the CRISPR-Cas9 system. The protein-RNA complex starts the search process from the solution, which is labeled as a state 0 (see Fig. 1). It can associate then to one of *l* PAM sites (type 1 on Fig. 1) on DNA with a rate *k*_*on*_. While bounded to the PAM sequence, which we also define as a PAM state, the RNA-protein complex has two possibilities. From the PAM state *i* (*i* = 1, 2*,…, l*), the RNA-protein complex has two possibilities. It might dissociate back into the solution with a rate *k*_*off*_, or it can start probing the DNA sequence next to the PAM by switching to the DNA interrogation state. The reversible transitions into these states (type 2 in Fig. 1) are described by rates *k*_1_ and *k*_2_, respectively. If the correct sequence is found(from the PAM state *i* = *m* in Fig. 1), the search process is accomplished. Otherwise, the Cas9-RNA molecule can start the process again by exploring other PAM sites after dissociating first into the solution, or it can cleave the DNA sequence with a rate *r* in the off-target process: see Fig. 1. It is important to note that the requirement for the initial binding to the PAM sequence is crucial for supporting the immune functions of CRISPR. It eliminates the possibility of self-targeting (i.e., cutting the chemically identical sequence on its own DNA) because the same sequences in the native genome are not flanked by the PAM segments(5).

**Figure 1:**
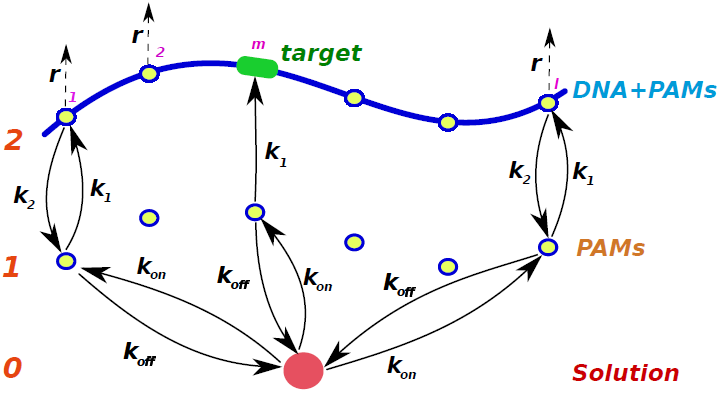
A discrete-state stochastic model to describe the DNA interrogation processes by CRISPR-Cas9 protein-RNA complexes. For convenience, a single Cas9-RNA molecule searching for the specific target on a single DNA molecule is considered. The DNA molecule of length *L* has *l* PAM sites, and one of them leads to the target sequence on DNA. The state 0 corresponds to the RNA-Cas9 complex in the solution. States of the type 1 (labeled also as PAM) correspond to the RNA-protein complex bound to the PAM site only, while the states of the type 2 (labeled also as DNA+PAM) describe the situation when the DNA interrogation is taking place. The association/dissociation rates from the solution to the PAM states are given by rates *k*_*on*_ and *k*_*off*_, respectively. The forward/backward transition rates between states of type 1 and 2 are given by *k*_1_ and *k*_2_, respectively. The rate for off-target cutting from the DNA+PAM states (not connected to the target) is equal to*r*.

**Table 1:** Prediction of DNA cleavage fraction based on equilibrium assumptions and comparison it with experimental values.

| $c_0, nM$ | $x, nM$ | $(c_0 - x), nM$ | $\frac{c_0 - x}{25} \times 100\%$ | Exp. |
| --- | --- | --- | --- | --- |
| 2.5 | 0.054 | 2.45 | 10% | 11% |
| 5 | 0.12 | 4.8 | 20% | 20% |
| 12.5 | 0.46 | 12 | 48% | 40% |
| 25 | 3.29 | 19 | 86% | 66% |
| 50 | 25.48 | 24.51 | 98% | 93% |

It has been argued before, that the protein search for the specific sites can be associated with first-passage processes (19, 24, 29–31). Then we can introduce a function *F_i_^(a)^(t)*, which is defined as a probability density to reach the target for the first time at time *t* if at *t* = 0 the protein was at the state *i*^(a)^, where *a* = 0, 1, 2 describes the type of the state (the solution, bound to the PAM state or the DNA interrogation state): see Fig. 1. The temporal evolution of these probability functions can be described utilizing the backward master equations (19, 24, 29, 31),

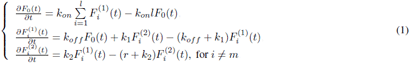

In addition, if the protein-RNA complex starts at *t* = 0 at the target site *m*^(2)^, the search process is instantly accomplished. This condition can be written as

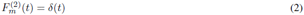

It is convenient to analyze the dynamics in the system using the method of Laplace transformations, when 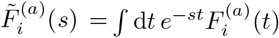. Then the original set of backward master equations is modified into a simpler system of algebraic equations,

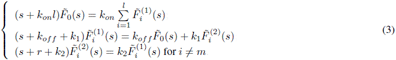

The initial condition Eq. (2) can now be written as

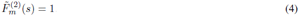

Obtaining exact expressions for the first-passage time probability density functions allows us to explicitly estimate all dynamic properties of the target search. For example, the overall probability of cutting the correct sequence, *P*_*c*_, and the mean search time to reach the correct sequence, *T*_0_ can be easily evaluated. In the first-passage language these quantities are associated with the splitting probability and the conditional mean first-passage time, which can be written as (24, 25)

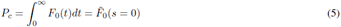

and

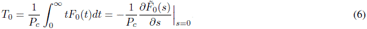

Solving Eqs. (3) leads to the following expression,

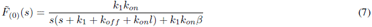

where we introduced an auxiliary function *β*,

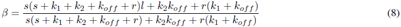

Then from Eqs. (5), (7) and (8) it can be shown that the probability of finding the correct target is given by

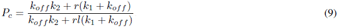

When there is no off-target cutting (*r* = 0), as expected, we have *P*_*c*_ = 1. The mean search time can be estimated using Eqs. (6), (7), (8) and (9), leading to

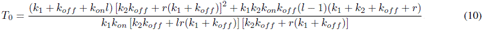

In the case of zero off-target cutting rates, the expression for the mean search time simplifies into

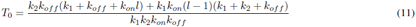

The physical meaning of this equation can be understood if we rewrite it as

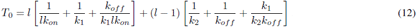

The first term corresponds to the time to go from the solution to the DNA interrogation state via the intermediate PAM state. On average, the protein-RNA complex will make *l* such attempts before the correct sequence is found. The second term describes the time it takes for the protein-RMA complex to return from the wrong sequence (*i* ≠ *m*) to the solution to start the search again. There are, on average, *l* − 1 such trajectories because the last excursion to the DNA interrogation state will be successful. Note also that the total rate out of the solution to PAM states is equal to *lk*_*on*_: see Fig. 1.

Our method can also calculate transient dynamic properties such as the probability to cleave DNA at time *t*,

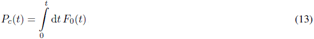

which is related to a time-dependent survival probability,

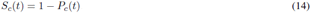

## Results and Discussion

### Explanation of Single-Turnover Observations

It is known that enzymes are biological catalysts that accelerate biochemical reactions without participating in them, and because of this each protein can act multiple times. Cas9 are helicase enzymes that stimulate the cleavage of DNA segments at specific locations. However, *in vitro* experimental measurements suggested that the behavior of the Cas9-RNA complexes deviates significantly from typical enzymes, which show multiple turnovers and follow Michaelis-Menten kinetics (5). It was found that the Cas9-RNA molecule can cut DNA, but it remains tightly bound to both after-cleavage nucleic acids segments at these experimental conditions (5). Only adding 7 M of urea allows Cas9-RNA to release the cleavage products. But it was also found, surprisingly, that the amount of cleaved DNA product was proportional to the molar ratio of Cas9-RNA and target DNA molecules (5). In other words, it means that the Cas9-RNA molecule effectively participates in the reaction and it is not a catalyst anymore!

To explain these surprising observations, we suggest that at these *in vitro* conditions the CRISPR system reaches the equilibrium with respect to association/dissociation of the Cas9-RNA complex and DNA. The protein finds to the target sequence on DNA (via the intermediate MAP states as indicated in Fig. 1), and it might cut the DNA. But because the cleavage products stay together, they can reverse the cleavage reaction: recall that all chemical reactions are reversible. The Cas9-RNA complex can eventually dissociate back into the solution. It is assumed here that the protein-RNA complex is tightly bound to DNA only when the DNA segment is cut. When the cut is healed the protein can dissociate back into the solution (probably again via the MAP state). This sequence of events essentially describes the binding/unbinding equilibrium in the system, which can be written as

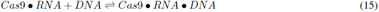

The dissociation equilibrium constant for this chemical reaction is

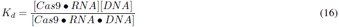

where [*Cas*9•*RNA*], [*DNA*] and [*Cas*9•*RNA*•*DNA*] are equilibrium molar concentrations of free Cas9-RNA complexes, free DNA molecules and Cas9-RNA complexes bound to DNA, respectively.

Equilibrium dissociation constant has been measured in experiments, yielding *K*_*d*_ ≃0.5 nM, and the starting concentrations of free Cas9-RNA complexes are also known (5). We define them as *c*_0_. Let us assume that at equilibrium we have [*Cas*9•*RNA*]_*eq*_ = *x* nM. Then, the mass balance requires that [*Cas*9•*RNA*•*DNA*]_*eq*_ = (*c*_0_ -*x*) nM, and [*DNA*]_*eq*_ = [25 − (*c*_0_ -*x*)] nM, because the initial concentration of DNA was 25 nM in these experiments (5). Substituting these equilibrium values into Eq. (16), we obtain,

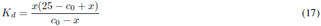

which allows us to explicitly evaluate the concentration (in nM) for every given value of *c*_0_ and *K*_*d*_. For experimentally measured *K*_*d*_ = 0.5 nM, one can derive,

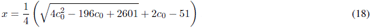

The results of these equilibrium calculations for different values of initial concentrations of Cas9-RNA complexes, *c*_0_, are presented in Tab. 1 and in Fig. 2, where they are also compared with experimentally measured values (5). It shows that this *in vitro* CRISPR system reaches the equilibrium already after several minutes. Note that the fraction of cleaved nucleicacids never reaches 100%, even when the initial concentration of Cas9-RNA is much larger than the stoichiometric ratio 1: 1 suggested by Eq.(15). This observation clearly supports the idea that the system goes into the equilibrium state. Thus, our simple equilibrium arguments capture quite well experimental observations, and the deviations could be explained by large errors bars in experimental measurements as well as by the possibility that the enzymes were not fully active at these experimental conditions (5). In addition, our theoretical explanations suggest that *in vivo* CRISPR systems have some additional biochemical components that allow the Cas9-RNA complex to be released after cutting the DNA chain.

**Figure 2:**
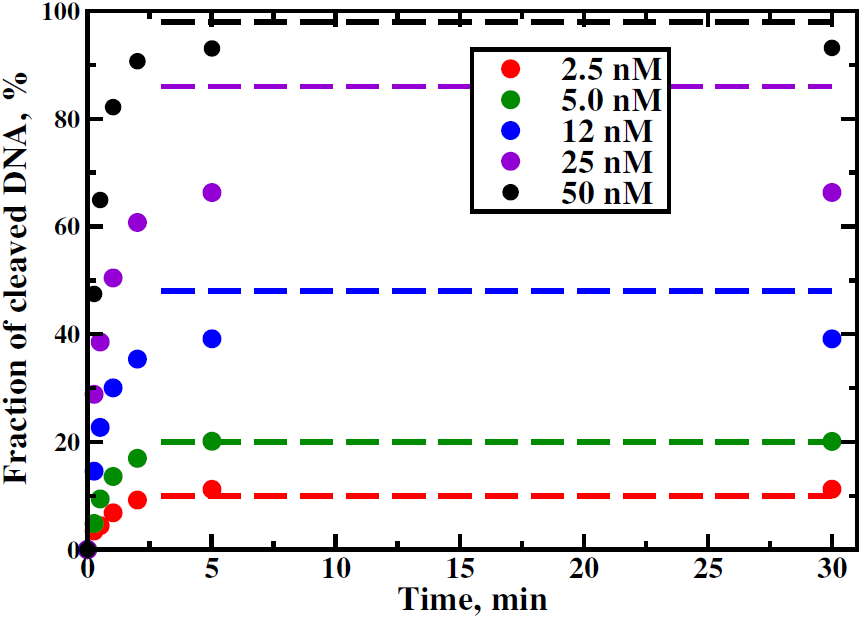
Comparison between experimental measurements (filled circles) and theoretical predictions (dashed lines) for the fraction of cleaved DNA as a function of time. The experimental data are taken from Ref.(5).

### Dynamics of Search without Off-Target Cutting

Now let us consider dynamics of target search by RNA driven Cas9 proteins, assuming a simpler situation when there is no off-target cutting. This corresponds to *r* = 0: see Fig. 1. Using the analysis presented above, we can compute the mean search times for Cas9-RNA complexes to locate specific sequences on DNA for different sets of parameters. The results are presented in Fig. 3, 4 and 5 for a realistic set of kinetic rates.

**Figure 3:**
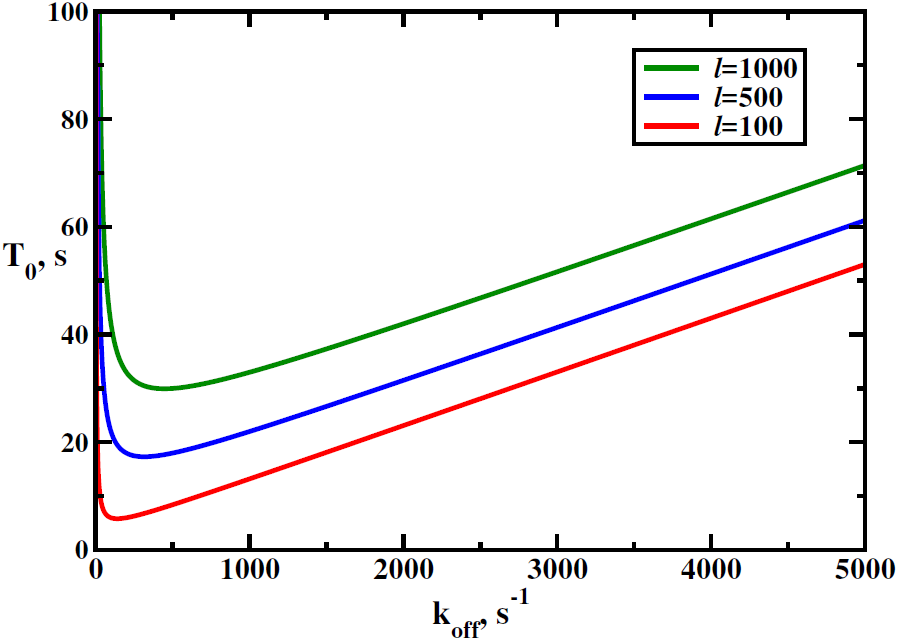
Mean search times as a function of the dissociation rate from the MAP, *k*_*off*_, for different numbers of PAMs, *l* [see Eq. (12)]. Parameters used for calculations are: *k*_1_ = *k*_2_ = 100 s^*-*1^and *k*_*on*_ = 1 s^*-*1^.

**Figure 4:**
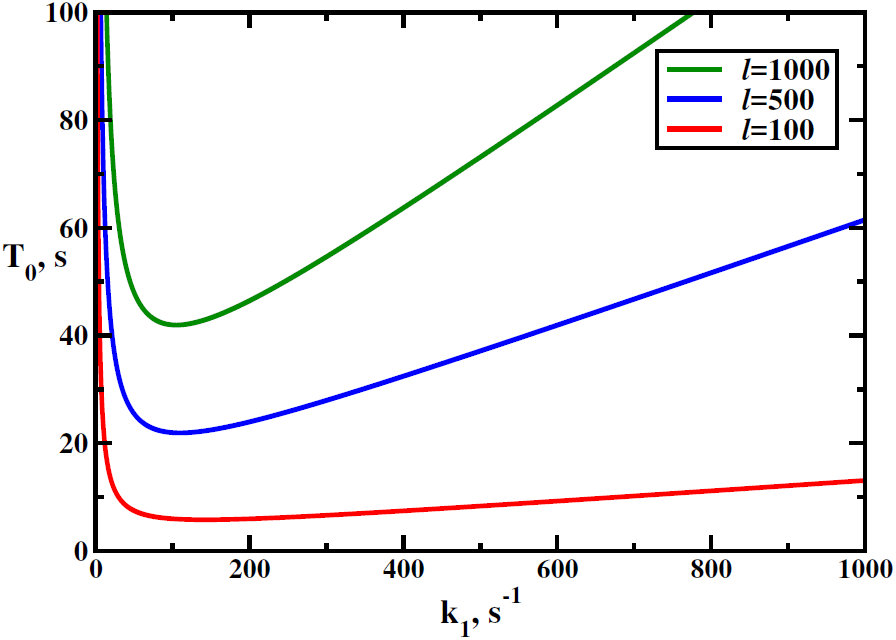
Mean search times as a function of the transition rate into the DNA interrogation state, *k*_1_, for different numbers of PAMs, *l* (see Eq. (12)). Parameters used for calculations are: *k*_*off*_ = *k*_2_ = 100 s^*-*1^and *k*_*on*_ = 1 s^*-*1^.

**Figure 5:**
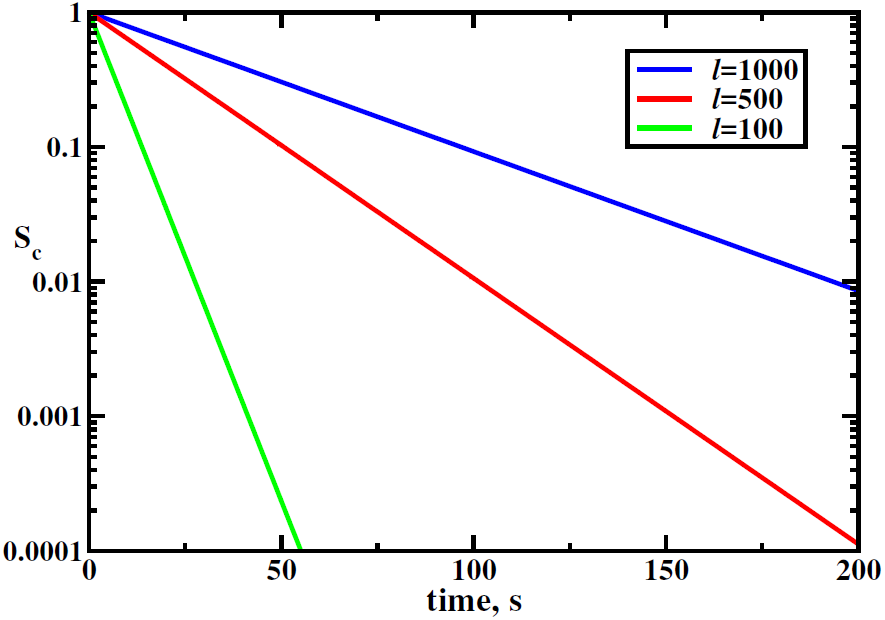
Survival probability for the DNA molecule to remain intact as a function of time. Parameters used for calculations are: *k*_1_ = *k*_2_ = *k*_*off*_ = 100 s^*-*1^, *k*_*on*_ = 1 s^*-*1^, and *l* = 2400.

Fig 3 shows the variation of the mean search time as a function of the dissociation rate *k*_*off*_ (see Fig. 1) from the MAP state back into the solution for different number of MAP states. It s found that *T*_0_ depends non-monotonically on the dissociation rate. These observations can be explained using the following arguments. For *k*_*off*_ ≪1 the protein-RNA complex might be trapped at the off-target locations along the DNA chain, preventing it from finding quickly the correct DNA sequence. For large dissociation rates, the situation is different: the Cas9-RNA molecule cannot go through the MAP state into the DNA interrogation state, and this also slows down the search dynamics. Fig. 3 also shows that increasing the number of MAP states slows down the search time the protein now must, on average, scan more sites, leading to larger *T*_0_.

Fig. 4 analyzes the mean search times as a function of the transition rate *k*_1_ from the MAP state into the DNA interrogation state: see Fig. 1. Again, a non-monotonic behavior is predicted in our model. For small *k*_1_, the Cas9-RNA complex cannot switch fast into the DNA interrogation state, significantly slowing the overall search dynamics. For fast transition rates(*k*_1_≪ 1), the protein-RNA complex will be trapped for longer periods of time at the off-target DNA interrogation states, and this also increases *T*_0_. One can see that this trapping effect is relatively weak for small number of MAPs, while increasing*l* slows down the search dynamics much stronger. This is expected because the larger *l*, the more traps for the Cas9-RNA complex in the system.

It is interesting to note that both rates, *k*_*off*_ and *k*_1_ can be associated with the strength of interaction between protein-RNA complex and MAP sequence on DNA. The non-monotonic character of the search dynamics suggests that there is an optimal strength of interactions. From this point of view, one could speculate that the relative short size of MAP sequences (three nucleotides) might be related with this observation. Longer MAP sequences would lead to stronger interactions, which might slow down the search, while shorter sequences would correspond to weaker interactions and larger number of PAMs per each DNA molecule.

Fig. 5 presents the survival probability of the DNA molecule not to be cleaved by the Cas9-RNA complex for different number of MAPs. One can see that the survival is higher for large *l*, as expected, since the helicase will take longer to locate and cut the correct DNA sequence. Theoretical results are similar to experimental observations reported in Ref. (5), further supporting our theoretical ideas.

### Dynamics of Search with Off-Target Cutting

Experiments suggest that the CRISPR-associated proteins Cas9 sometimes can cut the DNA sequences outside of the target segment (6, 11). It was argued that this might be the way for bacteria to fight quickly mutating viruses, in which the original target sequence would slightly change (17, 28). The probability of such off-target cutting events can reach up to 10-20%, and this significantly complicates the application of CRISPR systems for genetic applications (11). Our theoretical approach canquantitatively takes into account the possibility of the off-target cleavage of nucleic acids. The results of calculations using the full discrete-state stochastic model with in Fig. 1 with *r* ≠ 0 are presented in Figs. 6 and 7.

**Figure 6:**
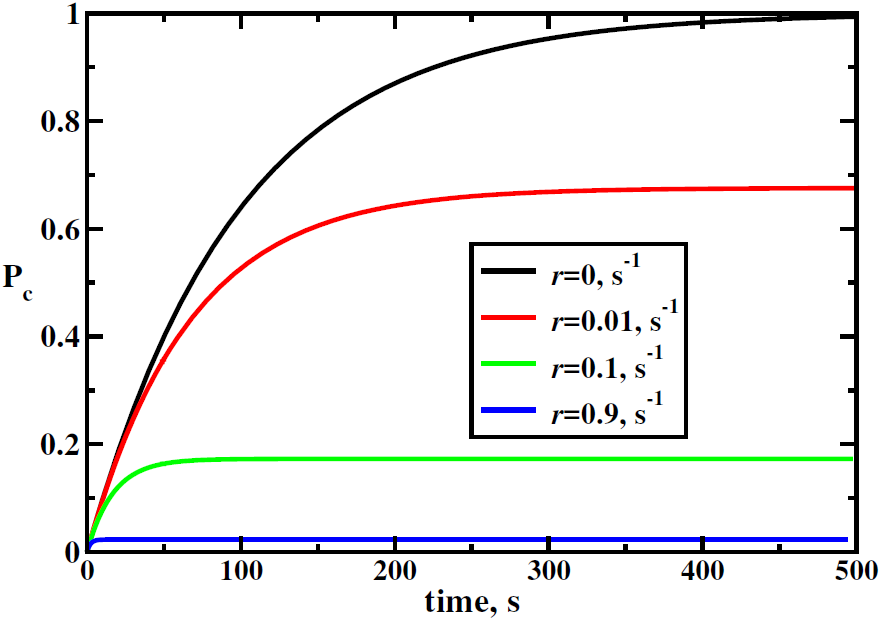
Probability of cutting DNA as a function of time for different values of the off-target rates *r* [see Eqs. (7) and (13)]. Parameters used for calculations are: *k*_1_ = *k*_2_ = *k*_*off*_ = 100 s^*-*1^, *k*_*on*_ = 1 s^*-*1^, and *l* = 2400.

**Figure 7:**
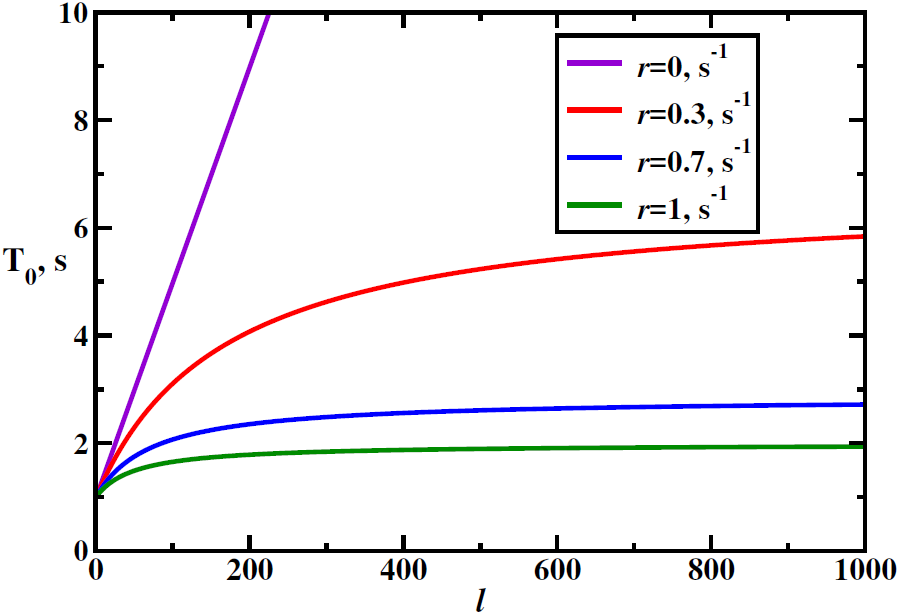
Mean search time as a function of the number of PAMs [see Eq. (10)]. Parameters used for calculations are:*k*_1_ = *k*_2_ = *k*_*off*_ = 100 s^*-*1^and *k*_*on*_ = 1 s^*-*1^.

Fig. 6 illustrates how the probability of target sequence cleavage, *P*_*c*_, changes with time for different off-target cutting rates *r*. In all cases, the probability first increases until it saturates to a constant stationary-state value. Increasing the off-target rate *r*, obviously, lowers this probability, but it also has another interesting effect. For larger *r* the CRSPR system reaches the stationary-state faster: for the parameters utilized for our calculations, it takes several hundreds seconds to achieve the steady-state conditions for *r* = 0, while for *r* = 0.9 s*-*^1^the stationary state is reached within a few seconds: see Fig. 6. It is easy to explain such behavior. The stationary-state probability of cleavage is lower for high off-cutting rates *r* and it is larger for smaller *r*. Thus, the stationary state can be reached much faster from the initial cleavage probability (*P*_*c*_ = 0 at *t* = 0) for fast off-cutting rates.

It is also interesting to analyze the mean search times in the more realistic CRISPR system with the off-target cutting, as presented in Fig. 7. The most surprising result here is that for the fixed set of parameters increasing the off-target rate *r accelerates* the search and decreases the mean search time *T*_0_. This can be understood if we recall that the mean search time is computed in our model as a conditional mean first-passage time to reach the target. So it is the time calculated by averaging the search times only over successful trajectories that lead to the cleavage. Increasing the off-cutting rate *r*, lowers the fraction of such trajectories and only the fastest trajectories that lead the protein-RNA complex faster to the target survive. The longer the Cas9-RNA molecule stays in the system, the higher is the probability to be removed via the off-target cutting. It is interesting to speculate if this effect has a biological significance. One could suggest that bacteria might intentionally tune the off-target rate *r* in the CRISPR immune response to increase the speed of locating and eliminating the foreign nucleic acids, sacrificing at the same time the accuracy of the process. Probably, cuts made at wrong locations of its DNA can be also healed via the correcting mechanisms, while the cleavage of foreign DNA segments is irreparable. It will be interesting to explore this idea further.

Fig. 7 also shows the dependence of the search time on the number of MAPs in the system. The smallest time is found for *l* = 1, but it always take longer to search for larger *l*, although the effect decreases significantly for high values of the off-target cutting rates *r*. This is because, as we explained above, at these conditions only the fastest trajectories leading to the target sequence survive, and this is not affected much on the total number of MAPs.

## Conclusions

We constructed a minimalist computational model for the search of specific target sequences on DNA by CRISPR-associated Cas9 protein-RNA complexes. It is based on the idea that the search is taking place via two step process: first the Cas9RNA complex attaches to the special tri-nucleotide PAM sequence on DNA, and then it can reversibly transition into the DNA interrogation state where the complementarity between RNA and DNA segments is utilized to recognize the specific target sequence. A discrete-state stochastic model of the Cas9-RNA target search, which takes into account the most relevant physical-chemical processes in the system, is developed. It is solved analytically using the method of first-passage processes, providing a comprehensive description of the dynamic processes in the CRISPR system. Our theoretical method is employed then to understand experimental observations in the CRISPR-Cas9 system. We propose and quantitatively test the idea that the single-turnover observations for Cas9 enzymes describe the effective equilibrium between free and DNA-bound Cas9-RNA molecules. In the next step, the search dynamics by RNA0guided Cas9 proteins is analyzed for the simplified case of no off-target cutting. It is found that the mean search times behave non-monotonically as the function of the dissociation and transition rates out the state where the Cas9-RNA complex is bound to the MAP sequence. These observations are explained by arguing that the protein-RNA complex can be trapped in off-target DNA interrogation states, or the search can be not fast due to slow passing the intermediate MAP states. It is argued that minimal search times correspond to the optimal interaction strength between Cas9-RNA and MAP sequences. A more realistic analysis with the possibility of off-target cleavage of the nucleic acids produces more interesting results. Our calculations show that the system reaches the stationary state with the probability of reaching the target sequence lower for fast off-cutting rates. The relaxation rate to the steady-state conditions is found to be faster for the higher off-cutting rates, which is explained using the first-passage arguments. It is also found that the mean search times decrease with more frequent off-target cleavages, and the significance of this finding for biological systems is discussed. It is proposed that bacteria might utilize this feature to accelerate the search and elimination of the foreign DNA material. Furthermore, the effect of the number of MAP states on the search dynamics is also analyzed.

Although our theoretical method is able to explain some experimental observations, it is important to critically evaluate it. Many features of the CRISPR systems are not taking into account in the proposed model. They include: 1) the neglect of the sequence dependence for utilized kinetic rates, which must depend on the chemical composition of the MAP sites, as well as on the chemical nature of the off-target sequences; 2) the oversimplification of the DNA interrogation process, which must involve multiple intermediate states when the RNA segment is trying to recognize the complementary DNA segment; and 3) the neglect of the cellular structure and crowding, which might affect these processes. For example, it is known that the target search is dependent on the local chromatin environment in live cells (6). However, despite these shortcomings, our theoretical model is very simple and it is able to capture main physical-chemical features of the target search by CRISPR associated Cas9-RNA complexes. The main advantage of our theoretical approach is that it provides a fully analytical description of dynamic properties, and it gives quantitative experimentally verifiable predictions. Obviously, it will be important to test our theoretical predictions using more advanced theoretical and experimental methods.

## Acknowledgments

A.B.K. acknowledges the support from the Welch Foundation (grant No. C-1559), from the NSF (grant No. CHE-1360979), and from the Center for Theoretical Biological Physics sponsored by the NSF (grant No. PHY-1427654).

